# MicrobiomeKG: Bridging Microbiome Research and Host Health Through Knowledge Graphs

**DOI:** 10.1101/2024.10.10.617697

**Authors:** Skye L. Goetz, Amy K. Glen, Gwênlyn Glusman

**Author notes:** Correspondence, Gwênlyn Glusman, Institute for Systems Biology, 401 Terry Ave N, Seattle, WA 98109, USA.

## Abstract

The microbiome represents a complex community of trillions of microorganisms residing in various body parts, and plays critical roles in maintaining host health and well-being. Understanding the interactions between microbiota and host offers valuable insights into potential strategies to promote health, including microbiome-targeted interventions. We have created MicrobiomeKG, a Knowledge Graph for microbiome research, bridging various taxa and microbial pathways with host health. This novel knowledge graph derives algorithmically-generated knowledge assertions from the supplemental tables supporting published microbiome papers, and is deployed for integrative analysis and hypothesis generation via the Biomedical Data Translator ecosystem.

## Introduction

The microbiome represents a complex community of trillions of microorganisms residing in various body parts, and plays critical roles in maintaining host health and well-being. Emerging research reveals that the microbiome influences numerous physiological processes, including digestion [1], aging [2], and immune system function [3]. Conversely, dysregulation of microbiota (dysbiosis) is associated with various diseases and negative health outcomes (e.g., inflammatory bowel disease, obesity, diabetes, and neurological disorders) [1]. Hence, understanding the interactions between microbiota and host offers valuable insights into potential strategies to promote health, including microbiome-targeted interventions.

The NCATS Biomedical Data Translator (‘Translator’) is a cutting-edge platform aiming to revolutionize biomedical research [4]. It integrates vast amounts of diverse data, from genes to clinical records, and uses advanced algorithms to uncover insights and accelerate discoveries. Harmonizing data and enabling semantic searches, Translator fosters collaboration among researchers and facilitates the development of new treatments and therapies for various diseases. The Translator project uses Knowledge Graphs (KGs) to store the wealth of data required to reason in a compact, easy-to-parse, universal format. KGs organize data from multiple sources, capture information about entities of interest in a given domain or task, and display connections between them. KGs comprise nodes (things) and edges (relationships between things).

Prominent projects come close to reconciling microbiome data with Translator philosophy. BugSigDB [5] serves as a comprehensive database of published microbial signatures, but lacks content connecting the microbiome and host health, and a Knowledge Graph format.

KG-Microbe [6] is an integratively analyzable Knowledge Graph linking prokaryotic data for phenotypic traits, taxonomy, chemicals, and environment descriptors, but is yet to include content linking the microbiome and host health. MicroPhenoDB [7] incorporates content linking the microbiome with host health, but lacks a Knowledge Graph format. MetagenomicsKG [8] incorporates multiple content sources, inclusive of microbiome-host health knowledge, into an integratively analyzable Knowledge Graph. However, neither includes knowledge from supplemental tables in their findings.

Here we present MicrobiomeKG, a integratively analyzable Knowledge Graph for microbiome research bridging various taxa and microbial pathways with host health, built from algorithmically-generated knowledge assertions from supplemental tables, and deployed to Translator [4].

## Methods

### Selection of Publications and Supplemental Tables

The publications included in the initial version of MicrobiomeKG represent a manually curated set of recent and multiomic-driven scientific papers that (a) bridge microbiome and host health-related content, and (b) include one or more supplemental tables with content that can be modeled as subject-predicate-object triples (e.g., taxon X affects disease Y) - the standard units of Knowledge Graphs.

### Derivation of Knowledge Assertions

Leveraging relevant content from the supplemental tables, their descriptions, or the manuscript itself, we ingest supplemental data contents into DataFrames using Python’s “polars” library [9]; edge attributes are manually computed when not provided but reasonably inferred (e.g., the total cohort size for meta analyses where the cohort sizes for all initial analyses are made explicit). We use custom Python scripts to transform the DataFrames values in multiple ways, via operations on individual values and on entire rows. Value transformations include: mathematical transformations (e.g., exponentiating log-transformed p-values), extracting relevant content with Python’s “re” library substrings (e.g., extracting “Actinobacteria” from “kurilshikov_class.Actinobacteria.id.419”), and text cleaning (e.g., deriving “enterocloster bolteae” from “enterocloster_bolteae”). Row operations include: filtering based on certain conditions (e.g., based on a given column’s Boolean value), dropping duplicates, dropping null values, and imposing cutoffs for filtering. We use a p-value cutoff of 0.075, thus the graph contains both statistically significant, and not significant but highly suggestive edges.

### Standardization of KG Contents and Structure

We standardize all edge predicates and node categories to Biolink ontology predicates and Biolink ontology classes [10]. Further, we map nodes to ontologies, representing them using compact universal resource identifiers (CURIEs), and normalize them using BABEL (version of 08/14/2024) [11]. We drop any knowledge assertions that fail to map subject or object to standard CURIEs. Finally, we export the output as Knowledge Graph Exchange (KGX) tab-separated values (TSV) format [12].

### Deployment

We deploy MicrobiomeKG as a public web application programming interface (API) that uses Translator Reasoner API (TRAPI) [13] format. We achieve this using Plover [14], an in-memory Python-based platform designed to host and serve Biolink-compliant knowledge graphs as TRAPI APIs. Plover enables one-hop queries of the underlying KG and automatically performs Biolink predicate/class hierarchical reasoning and concept subclass transitive chaining, among other tasks. The Plover MicrobiomeKG API is accessible at multiomics.test.transltr.io/mbkp.

## Results

### Overview of MicrobiomeKG

We developed Microbiome KG, a Knowledge Graph built for microbiome research, focusing on the interface between the microbiome and the health of the host. The project contains knowledge assertions crafted from 66 different supplemental tables (listed in Supp. Table 1) across 22 publications. The number of assertions derived from each publication varies over three orders of magnitude (Fig. 1), reflecting the huge diversity in content and level of detail of supplemental tables. The KG comprises 5,365 nodes (concepts) and 45,133 edges (assertions) that outline relationships between the microbiome and various host health factors, spanning 28 Biolink [10] ontology classes (most commonly genes, taxa, and chemicals, Fig. 2) and four Biolink ontology predicates (most commonly representing associations and correlations, Fig. 3).

**Fig. 1:**
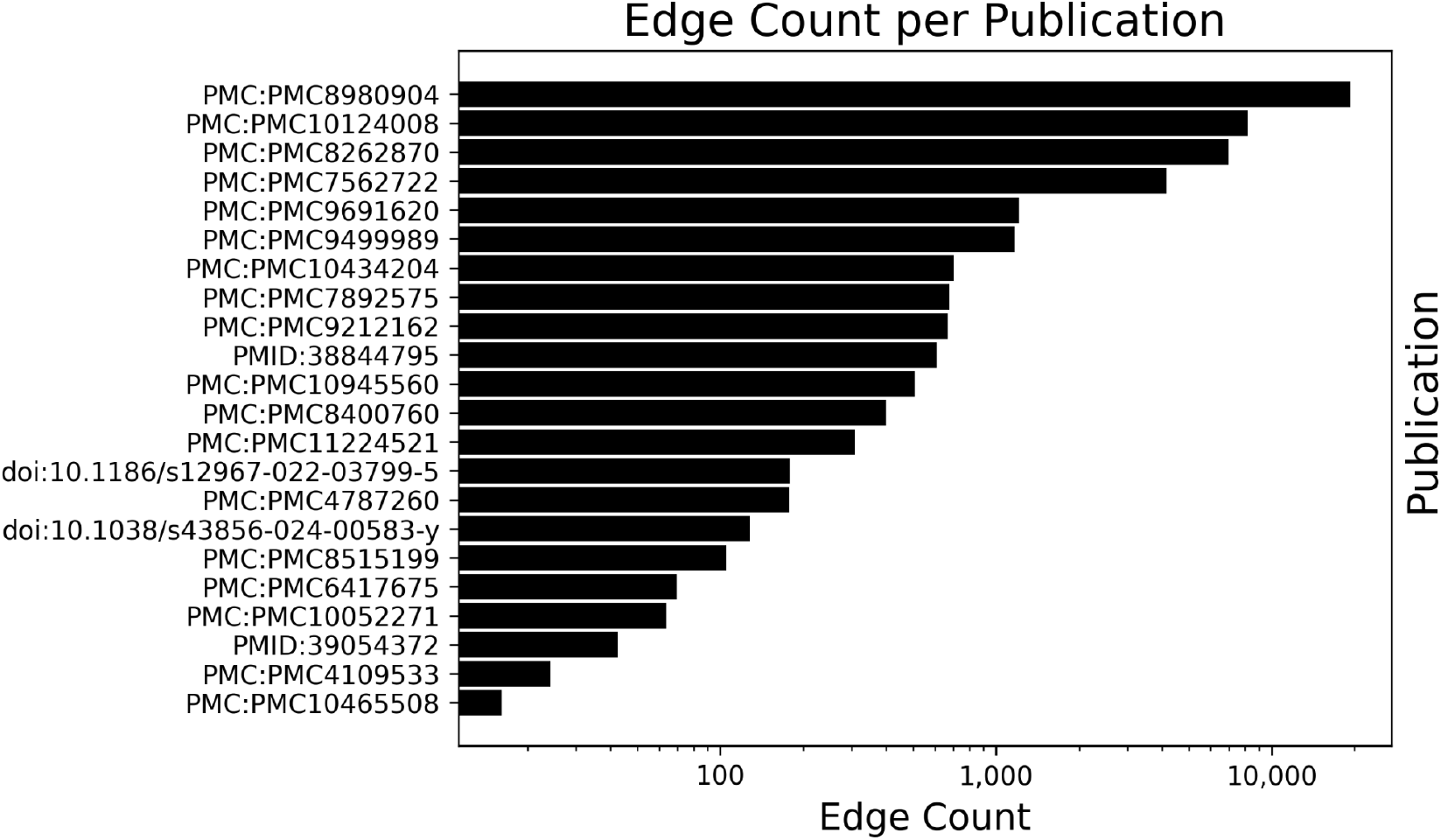
Edge Count per Publication. Number of edges in MicrobiomeKG per included publication, sorted from most to least frequent.

**Fig. 2:**
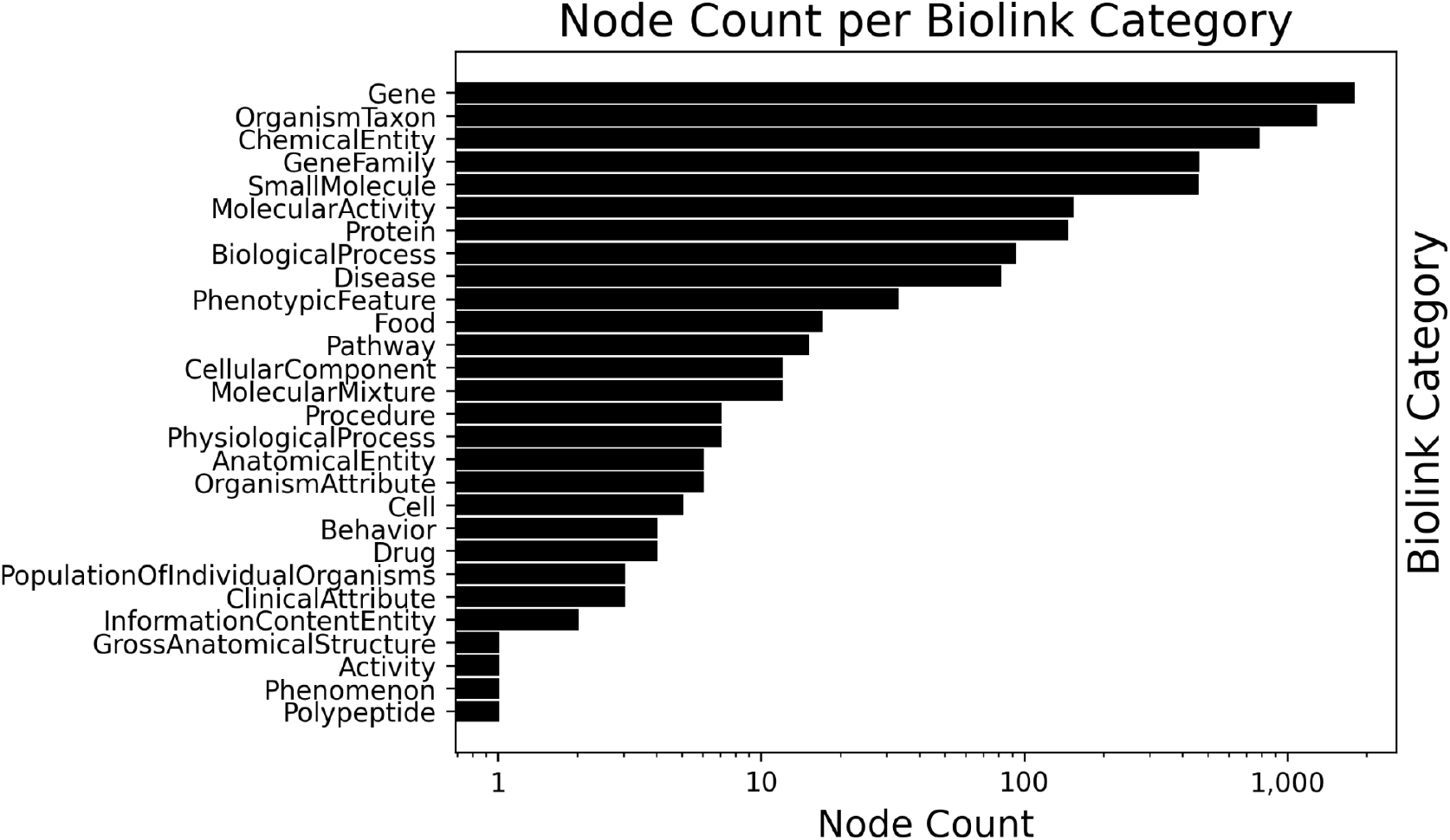
Node Count per Biolink Category. Number of nodes in MicrobiomeKG per Biolink ontology class, sorted from most to least frequent.

**Fig. 3:**
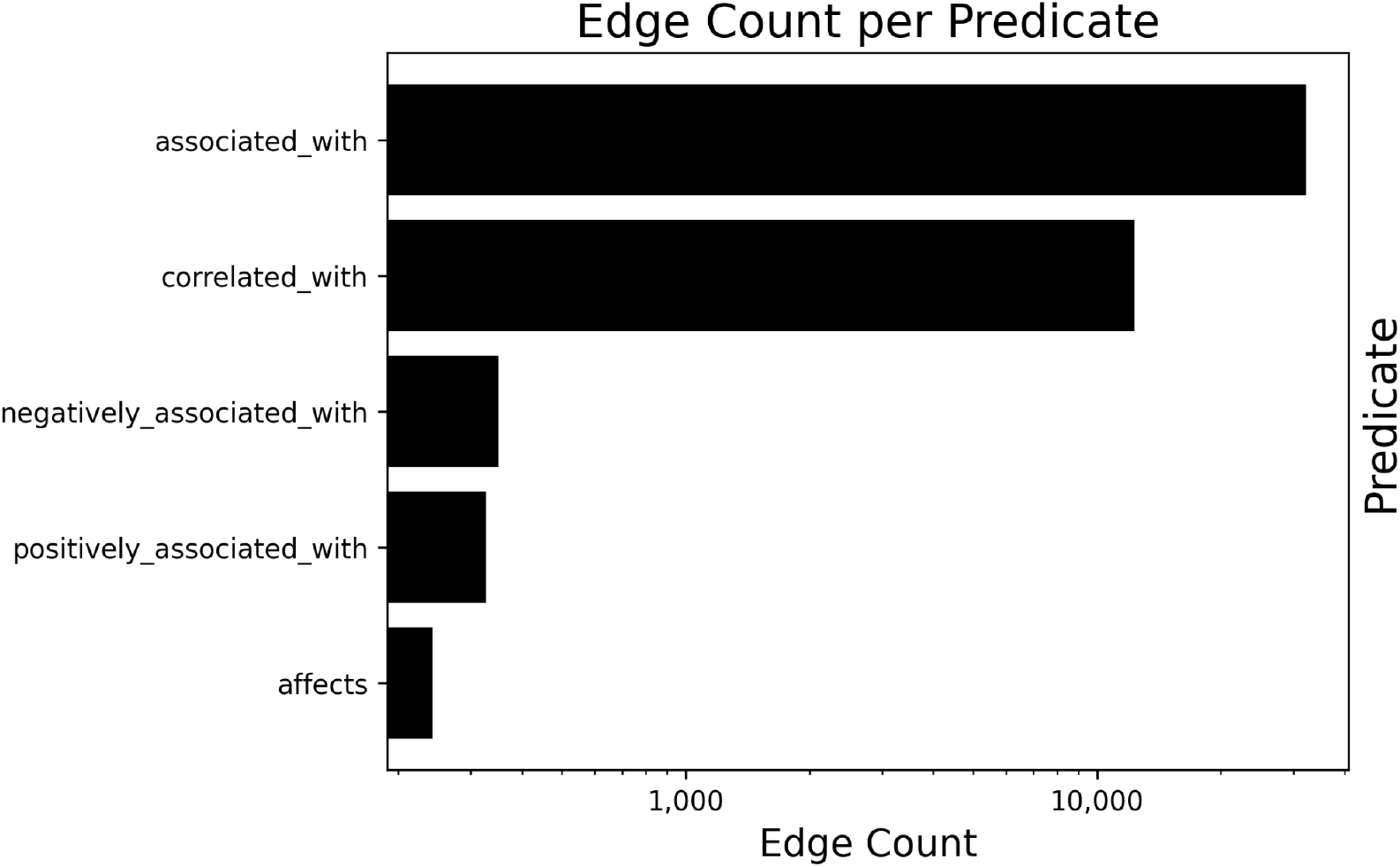
Edge Count per Predicate. Number of edges in MicrobiomeKG per Biolink ontology predicate, sorted from most to least frequent.

### NAFLD/Helminthiasis Case Study

Through the combination of edges derived from two publications already integrated into MicrobiomeKG, we identified a hypothetical connection between helminthiasis (MONDO:0004664) and metabolic dysfunction-associated steatotic liver disease (also known as non-alcoholic fatty-liver disease, or NAFLD, MONDO:0013209). This connection is consistent with and supported by published observations [15–17].

Helminthiasis is a global health burden, particularly in economically underdeveloped regions. Helminth colonization has been linked to changes in the host gut microbiome, namely increased diversity [18]. More recent work identified significant alterations in host gut and saliva microbiota, driven by clinical helminth infections [19], at multiple taxonomic levels. We highlight here (Fig. 4) the statistically significant association between helminth infections and gut bacteria of the orders Burkholderiales (adjusted p-value = 0.0026) and Lactobacillales (adjusted p-value ∼ 0), as reported in Supplemental Table 3 of Gobert *et al*., 2022 [19].

**Fig. 4:**
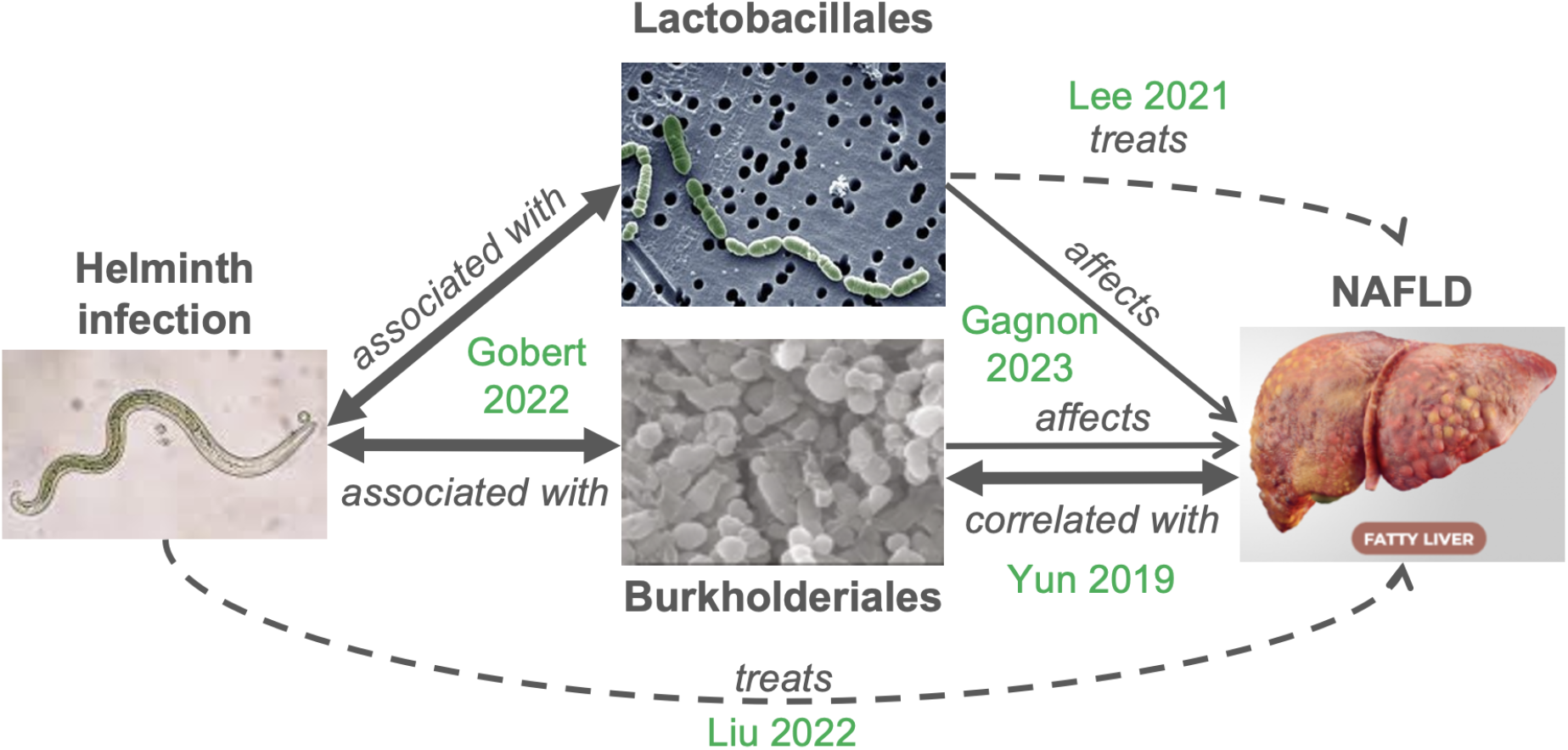
Microbiome Effects Link Helminth Infections with Non-Alcoholic Fatty Liver Disease (NAFLD). Arrows indicate relationships between the concepts (helminth infection, bacterial classes, NAFLD). Bidirectional arrows represent symmetric relationships (associated with, correlated with). Thicker lines represent significant relationships, dashed arrows represent higher-level ‘treats’ relationships. The source publication for each edge is indicated in green.

Non-alcoholic fatty liver disease (NAFLD) is a highly prevalent form of progressive and chronic liver disease, with gradual accumulation of liver fibrosis and cirrhosis. The pathogenesis of NAFLD is complex and involves disrupted glycolipid metabolism, inflammation, and dysregulation of the gut microbiota [20]. Metagenomic studies identified bacterial taxa positively or negatively associated with progression to advanced fibrosis in NAFLD [21]. Furthermore, Gagnon *et al*., 2023 used Mendelian randomization to establish the causal relationships between gut microbiota and multiple cardiometabolic traits and chronic diseases, including NAFLD [22]. We highlight their finding that Class Betaproteobacteria affects (leads to) NAFLD, with a Benjamini-Hochberg adjusted p-value of < 0.0218 as computed using the inverse variance weighted (IVW) method (their Suppl. Table 5) and adjusted p-value of < 0.000076 as calculated using the IVW radial method (their Suppl. Table 6). The association with Order Burkholderiales, within Class Betaproteobacteria, did not reach the significance threshold but was suggestive, with an adjusted p-value of < 0.075 (their Supp. Table 5). A significant negative connection between Burkholderiales (specifically, *Parasutterella*) and NAFLD was reported by Yun *et al*., 2019 [23]. Similarly, the relationship with Lactobacillus did not reach statistical significance (adjusted p-value < 0.0664), but a mechanistic relationship is reported elsewhere [16]. Both Burkholderiales and Lactobacillales have potential application as therapeutics for NAFLD [16,17].

## Discussion

We have presented MicrobiomeKG, a novel Knowledge Graph connecting the microbiome and host health, and a case study highlighting its application. The initial version of the graph contains 5,365 nodes and 45,133 edges, sourced from a set of 22 microbiome papers (Fig. 1), in an integratively analyzable KGX TSV format [12], deployed via Plover [14] to the Translator ecosystem.

MicrobiomeKG derives knowledge assertions drawn from supplemental materials published together with microbiome papers. Unlike the standard application of natural language processing of paper abstracts and or full texts of papers, which is, perforce, limited to content the authors decided to discuss in the text (and, potentially, the main-text tables), content extraction from the supplemental tables may capture a significantly larger corpus of knowledge assertions not included in the manuscript for a variety of reasons, including considerations of statistical significance, space limitations, and decisions about focus of narrative. In some cases, the supplemental tables provide precise numerical values for content included in the manuscript narrative in a simplified or approximate form, or perhaps in graphical form in embedded figures, which pose additional data extraction challenges. By table-mining the supplemental materials, we are thus able to maximize knowledge extraction while minimizing reproduction errors. For example, most of the edges underlying the NAFLD/Helminthiasis Case Study are not in their papers’ main text, tables, or figures. Yet they’re readily derivable from the supplemental data tables. Supporting materials from publications have been used to extract gene sets [24]; here, we applied them to extract structured knowledge assertions.

Supplemental materials can be very difficult to use [25]. By identifying knowledge assertions from supplemental tables, and expressing them as knowledge graphs, we are casting this valuable content into a format that is ideal for hypothesis generation. MicrobiomeKG ultimately brings to Translator novel nodes and edges that foster previously unexplored connections between the microbiome and varied biomedical data. We expect that MicrobiomeKG will be the first of many Knowledge Graphs built from knowledge assertions derived from the trove of untapped supplemental tables. In the context of graph machine learning, such extended knowledge extraction will prove advantageous for training microbiome, biological, biomedical, and host health AI/ML models [26].

## Author Contributions

GG conceived of the study. GG, SLG, AKG designed and implemented the system. GG, SLG performed analyses. GG, SLG wrote the manuscript. All authors edited and approved its final version.

## Acknowledgements

This work was supported by the National Science Foundation (Award #2150265) and by the National Center for Advancing Translational Sciences, Biomedical Translator Program (Other Transaction Awards OT2TR003443 and OT2TR003428). Any opinions expressed in this document are those of the Translator community at large and do not necessarily reflect the views of NCATS, individual Translator team members, or affiliated organizations and institutions. We wish to thank Jared C. Roach for advice and discussion.

## Data Availability

The NCATS Biomedical Data Translator, and by extension MicrobiomeKG, is publically accessible and can be queried through the Translator UI (https://ui.transltr.io).

The Plover MicrobiomeKG API is accessible at

- Development version: https://multiomics.rtx.ai:9990/mbkp
- Deployed version: multiomics.test.transltr.io/mbkp

## Supplemental Materials

**Supplemental Table 1:** Index of Supplemental Tables Included.

## References

1. Hills RD, Pontefract BA, Mishcon HR, Black CA, Sutton SC, Theberge CR. Gut Microbiome: Profound Implications for Diet and Disease. Nutrients. 2019;11. doi:10.3390/nu11071613

2. Wilmanski T, Diener C, Rappaport N, Patwardhan S, Wiedrick J, Lapidus J, et al. Gut microbiome pattern reflects healthy ageing and predicts survival in humans. Nature metabolism. 2021;3. doi:10.1038/s42255-021-00348-0

3. Wiertsema SP, van Bergenhenegouwen J, Garssen J, Knippels LMJ. The Interplay between the Gut Microbiome and the Immune System in the Context of Infectious Diseases throughout Life and the Role of Nutrition in Optimizing Treatment Strategies. Nutrients. 2021;13. doi:10.3390/nu13030886

4. Fecho K, Thessen AE, Baranzini SE, Bizon C, Hadlock JJ, Huang S, et al. Progress toward a universal biomedical data translator. Clin Transl Sci. 2022;15. doi:10.1111/cts.13301

5. Geistlinger L, Mirzayi C, Zohra F, Azhar R, Elsafoury S, Grieve C, et al. BugSigDB captures patterns of differential abundance across a broad range of host-associated microbial signatures. Nat Biotechnol. 2023;42: 790–802.

6. GitHub - Knowledge-Graph-Hub/kg-microbe. In: GitHub [Internet]. [cited 10 Oct 2024]. Available: https://github.com/Knowledge-Graph-Hub/kg-microbe

7. Yao G, Zhang W, Yang M, Yang H, Wang J, Zhang H, et al. MicroPhenoDB Associates Metagenomic Data with Pathogenic Microbes, Microbial Core Genes, and Human Disease Phenotypes. Genomics Proteomics Bioinformatics. 2020;18: 760–772.

8. Ma C, Liu S, Koslicki D. MetagenomicKG: a knowledge graph for metagenomic applications. bioRxiv. 2024. doi:10.1101/2024.03.14.585056

9. DataFrames for the new era. [cited 18 Sep 2024]. Available: https://www.pola.rs/

10. Unni DR, Moxon SAT, Bada M, Brush M, Bruskiewich R, Caufield JH, et al. Biolink Model: A universal schema for knowledge graphs in clinical, biomedical, and translational science. Clin Transl Sci. 2022;15: 1848–1855.

11. GitHub - TranslatorSRI/Babel: Babel creates cliques of equivalent identifiers across many biomedical vocabularies. In: GitHub [Internet]. [cited 23 Sep 2024]. Available: https://github.com/TranslatorSRI/Babel

12. GitHub - biolink/kgx: KGX is a Python library for exchanging Knowledge Graphs. In: GitHub [Internet]. [cited 17 Sep 2024]. Available: https://github.com/biolink/kgx

13. GitHub - NCATSTranslator/ReasonerAPI: NCATS Biomedical Translator Reasoners Standard API. In: GitHub [Internet]. [cited 18 Sep 2024]. Available: https://github.com/NCATSTranslator/ReasonerAPI

14. GitHub - RTXteam/PloverDB at multiomics. In: GitHub [Internet]. [cited 18 Sep 2024]. Available: https://github.com/RTXteam/PloverDB

15. The gut-liver-kidney axis: Novel regulator of fatty liver associated chronic kidney disease. Pharmacol Res. 2020;152: 104617.

16. Lee NY, Shin MJ, Youn GS, Yoon SJ, Choi YR, Kim HS, et al. Lactobacillus attenuates progression of nonalcoholic fatty liver disease by lowering cholesterol and steatosis. Clin Mol Hepatol. 2021;27: 110–124.

17. Liu X, Jiang Y, Ye J, Wang X. Helminth infection and helminth-derived products: A novel therapeutic option for non-alcoholic fatty liver disease. Front Immunol. 2022;13: 999412.

18. Lee SC, Tang MS, Lim YAL, Choy SH, Kurtz ZD, Cox LM, et al. Helminth colonization is associated with increased diversity of the gut microbiota. PLoS Negl Trop Dis. 2014;8: e2880.

19. Gobert GN, Atkinson LE, Lokko A, Yoonuan T, Phuphisut O, Poodeepiyasawat A, et al. Clinical helminth infections alter host gut and saliva microbiota. PLoS Negl Trop Dis. 2022;16: e0010491.

20. Han H, Jiang Y, Wang M, Melaku M, Liu L, Zhao Y, et al. Intestinal dysbiosis in nonalcoholic fatty liver disease (NAFLD): focusing on the gut-liver axis. Crit Rev Food Sci Nutr. 2023;63. doi:10.1080/10408398.2021.1966738

21. Loomba R, Seguritan V, Li W, Long T, Klitgord N, Bhatt A, et al. Gut Microbiome-Based Metagenomic Signature for Non-invasive Detection of Advanced Fibrosis in Human Nonalcoholic Fatty Liver Disease. Cell Metab. 2017;25: 1054–1062.e5.

22. Gagnon E, Mitchell PL, Manikpurage HD, Abner E, Taba N, Esko T, et al. Impact of the gut microbiota and associated metabolites on cardiometabolic traits, chronic diseases and human longevity: a Mendelian randomization study. J Transl Med. 2023;21: 1–14.

23. Yun Y, Kim H-N, Lee E-J, Ryu S, Chang Y, Shin H, et al. Fecal and blood microbiota profiles and presence of nonalcoholic fatty liver disease in obese versus lean subjects. PLoS One. 2019;14: e0213692.

24. Clarke DJB, Marino GB, Deng EZ, Xie Z, Evangelista JE, Ma’ayan A. Rummagene: massive mining of gene sets from supporting materials of biomedical research publications. Communications Biology. 2024;7: 1–13.

25. Pop M, Salzberg SL. Use and mis-use of supplementary material in science publications. BMC Bioinformatics. 2015;16: 1–4.

26. Knowledge graphs as tools for explainable machine learning: A survey. Artif Intell. 2022;302: 103627.

